# Pioneering New Breeding Pathways with 3PaTec: Unlocking the Power of Polyspermy-Induced Triparentage for Sustainable Crop Development

**DOI:** 10.1101/2025.03.14.643266

**Authors:** Saurabh Joshi, Christian A. Beer, Yanbo Mao, T Thomas Baum, Dawit G. Tekleyohans, Constantin Bubenheim, Joakim Palovaara, Olaf Czarnecki, Thomas Nakel, Rita Groß-Hardt

## Abstract

Over the last few centuries, advancements in plant breeding have revolutionized agriculture, driving significant increases in global food production. Polyploidy, the increase in chromosome copies, can positively affect plant performance and is assumed to have played a critical role in the domestication of crop plants. Polyploidy is thought to be primarily caused by sperm that, due to meiotic aberrations, deliver unreduced chromosome sets. We have recently identified an alternative pathway to polyploidization by demonstrating that polyspermy, the fertilization of an egg cell by more than one sperm, occurs in planta and results in viable triploid plants. Capitalizing on a novel high-throughput polyspermy detection tool, we have shown that polyspermy involving two pollen donors can generate plants with three parents, one mother and two fathers. This 3PaTec technology not only speeds up breeding processes through an instant combination of beneficial traits from three parents; it also allows selective polyploidization of the egg cell, thereby bypassing the central cell-derived embryo-nourishing endosperm, a major hybridization barrier. Here, we further explore the genetic and developmental factors influencing polyspermy and show that the frequency of polyspermy and triparental plant formation varies among ecotypes and depends on pollen availability, suggesting that polyspermy is an adaptive trait. Additionally, we extend the application of 3PaTec to crops by successfully generating triparental sugar beet in-field using a wind pollination strategy. Our findings highlight the potential of 3PaTec for major crop plants. This innovative breeding technology does not rely on genetic engineering, requiring minimal technical expertise and infrastructure. As a result, it is highly accessible to a wide range of users, contributing to the democratization of plant breeding by empowering individuals from all backgrounds to collaborate and contribute to developing resilient and sustainable crops.

## Introduction

Over the last few centuries, modern agriculture has undergone a dramatic transformation, significantly increasing global food production^1^: Early trial-and-error farming methods became more systematic with Mendel’s discovery of the principles of heredity in the 19th century. The discovery of experimental mutagenesis in the early 20th century paved the way to artificially increase genetic variation through radiation and chemical mutagens^2^. The mid-20th century Green Revolution introduced high-yielding crops responsive to fertilizers and pesticides, boosting food production but also highlighting environmental issues including soil degradation and a loss of biodiversity. Later, molecular biology techniques like marker-assisted selection were integrated into plant breeding, allowing for the selection of desirable traits linked to genetic markers. The advent of genetic engineering enabled direct manipulation of plant DNA and the introduction of DNA and traits from unrelated species, while modern gene-editing technologies like CRISPR now allow targeted modifications of distinct genetic loci.

Plant breeding has also harnessed the ability of plants to cope or even strive from changes in the number of chromosome copies (ploidy). Polyploidy is a natural phenomenon considered to be a major driving force for plant evolution and most plant lineages have undergone at least one whole genome duplication^3,4^. Domesticated plants have gone through more polyploidy events than their wild relatives, likely due to higher genetic degrees of freedom providing these lineages with phenotypic novelties that enhanced their adaptability during crop improvement^5,6^, including beneficial traits such as larger organ size, increased heterozygosity, and hybrid vigor. Apart from an increase in genome copies, also the induction of haploidy has proven highly valuable for the stabilization of desired traits^7,8^: The combination of genetically different genomes, for example, crossing climate-adapted wild varieties with elite cultivars, generates heterozygosity as an undesired side-effect. By developing haploid offspring from one parent and doubling their chromosomes, fully homozygous plants are rapidly produced. While haploid-inducing technologies have been established in Brassicas and cereals, such as barley, maize, rice, rye and wheat, they are not available to all crops^9^.

We have previously shown that plants not only tolerate a reduction in the number of parents but also an increase, which becomes possible due to rare deviations from the canonical reproductive mode: The fertilization of the two female gametes, egg and central cell, is typically associated with a disintegration of the pollen tube-attracting synergids and the establishment of chemical and physical egg cell barriers^10-12^. The combination of the polytubey and egg cell block effectively reduces the chances of supernumerary sperm entering the egg cell; however, these mechanisms are not 100% efficient. We have established a high-throughput polypaternal breeding assay called HIPOD which identifies the genetic material of both fathers combined in a single egg cell, a process that necessitates polyspermy. Using this technique we have demonstrated that *in-planta* polyspermy is a rare but plausible scenario: The *Arabidopsis thaliana* variation Landsberg *erecta* gives rise to 0.012 % polyspermy-induced plants^13^. Given the high number of seeds generated by Arabidopsis plants, this suggests that, on average, each plant produces one or more polyspermy-induced polyploids under our growth conditions. The resulting plants are viable, produce more biomass and exhibit a fertility rate of over 30% giving rise to thousands of viable offspring^13^. In addition, triparental plants segregate not only aneuploids but also diploid and tetraploid offspring in the next generation, which highlights the great potential of polyspermy for generating neopolyploids. This previously unrecognized route to polyploidization also explains an unresolved conundrum: In the past, unreduced gametes that arise infrequently from meiotic aberrations during pollen development have been considered the primary mechanism for natural polyploidization^14-16^. However, an endosperm hybridization barrier, also called the triploid block, is activated upon the delivery of gametes from pollen donors with higher chromosome copies or from unreduced male gametes^17,18^, leading to seed abortion. By contrast, polyspermy can selectively polyploidize the egg cell thereby bypassing this barrier^10,19^.

The finding that plants can have three parents carries significant implications for plant breeding. The technology of three-parent crosses not only enables the immediate integration of beneficial traits from three parents but also provides a means for the combination of plants that could previously not be combined due to hybrid incompatibilities that are induced in the endosperm^10,19^. In the current study, we demonstrate that the genetic background of different ecotypes, the amount of pollen deposited to the stigmatic surface, and the proximity of an ovule to the stigma influence polyspermy-induced generation of triparental plants. Additionally, we demonstrate the practical application of three-parent crosses in crops by transitioning the technique from a laboratory setting to field production, successfully creating triparental sugar beet through a wind pollination strategy optimized for balanced paternal pollen density. This highlights the potential of three-parent crosses as a powerful breeding tool.

## Results and discussion

Polyspermy, a previously unrecognized natural pathway to polyploid plants, has only recently been discovered and remains challenging to study. This is due to the difficulty of observing sperm-egg fusion and because it is a rare event, as it is vastly outnumbered by regular monospermic fertilization. Given the evolutionary and breeding significance of polyspermy, we sought to gain deeper insights by investigating whether its frequency is an inherent constant in plants or can be influenced by external factors.

### Polyspermy-induced polyploidization positively correlates with the number of available pollen

In an initial experiment, we investigated whether the quantity of pollen available to a single flower influences the frequency of polyspermy-induced polyploidization. To assess this, we utilized the HIPOD assay, a previously established method specifically designed to detect triparental seedlings resulting from polyspermy-induced polyploidization^13^. The HIPOD assay leverages the GAL4/UAS two-component system, allowing only seedlings fertilized by sperm from two distinct pollen donors to survive herbicide selection: Pollen donor one ubiquitously expresses the yeast GAL4 transcription factor, which when combined in a single egg cell activates the GAL4-responsive UAS promoter and an associated fluorophore-tagged herbicide resistance conferring BAR gene.

We implemented two pollination methods. In the “saturated” pollination approach, we coated the stigmatic surface with pollen using paint brushes, achieving a pollen-to-ovule ratio higher than typically observed in a self-pollinating species like Arabidopsis^20^. In contrast, the “limited” pollination method involved carefully placing only a few pollen grains onto the stigma using a single eyelash, creating a low pollen-to-ovule ratio (<1) and thereby limiting pollen tube availability (Fig. 1 A,B). In total, we processed over 3,400 siliques under the limited pollination condition and over 1,700 siliques under the saturated condition The length of the siliques varied between the two methods, reflecting differences in seed numbers: siliques from the saturated pollination generated an average of 65 seeds, while those from the limited pollination produced around 27 seeds per silique, confirming the effectiveness of each pollination approach.

**Figure 1:**
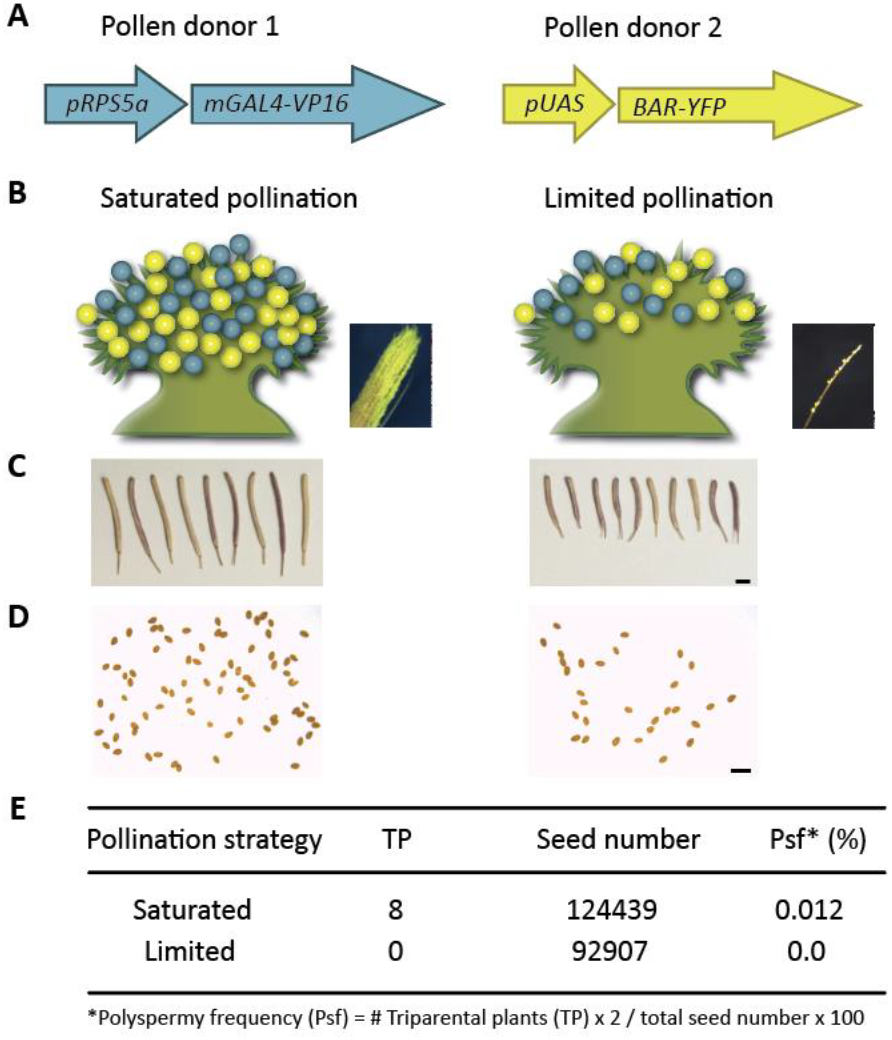
High pollen availability enhances the frequency of polyspermy-induced polyploidization. A) In the HIPOD assay, two pollen donors each encoding one component of a bipartite transactivating system provide herbicide resistance when combined via polyspermy in a single egg cell. B) Saturated and limited pollination strategies for the application of pollen from the two different pollen donors (blue and yellow) onto the stigmatic surface of a flower (green). Saturated pollination was performed (Movie 1) using a paintbrush (left inset), while limited pollination was conducted (Movie 2) with an eyelash (right inset; eyelash with a small number of pollen grains attached), ensuring only a few pollen grains were applied. C) Harvested siliques showing different lengths. D) Example of seeds obtained from a single silique subjected to either saturated (left) or limited pollination (right). E) Polyspermy frequencies obtained from both strategies. Scale bars 2 mm (C) and 1 mm (D). Figure adapted from Mao *et al*. 2020.

Crucially, eight triparental plants were obtained from the saturated pollination, while none were recovered from the limited pollination. This result highlights the importance of high pollen density for polyspermy and implies that polyspermy-induced triparental plant formation can be induced in densely populated environments where pollen competition is higher, offering valuable insights for optimizing breeding strategies.

### Positional effects: Polyspermy-induced triparental plants initiate in the distal portion of the gynoecium

Under our growth conditions, a typical silique of the Arabidopsis Col-0 accession aligns on average between 30 and 35 ovules along the distal proximal axis in the two locules of the gynoecium. Since the stigmatic surface is positioned at the distal end of the gynoecium, pollen tubes reach the most distal part of the gynoecium more quickly and in greater numbers compared to the proximal regions. Given that an increased number of pollen tubes enhances the likelihood of polyspermy, we hypothesized that ovules located more distally give polyspermy-induced triparental plants more often than those situated proximally. We have previously established an assay that allows the detection of polyspermy-derived triparental embryos already in the seed (HIPOD_SCO1_)^19^. This adaptation of the HIPOD assay exploits that homozygous mutants for the gene *SNOWY COTYLEDON 1* (*SCO1*) exhibit pale green seeds that can easily be distinguished from the darker green counterparts^21^. HIPOD_SCO1_ consists of two pollen donors that each contain one part of the yeast GAL4-UAS two-component system (*sco1/-pRPS5a:mGAL4-VP16/+* and *sco1/-pUAS::SCO1_tdTOMATO/+*), which when combined in an egg cell activate an intact *SCO1* copy in otherwise *sco1* homozygous pollen acceptor. Triparental embryos can then be detected early within siliques based on their darker seed colour^19^ (Figure 2A).

**Figure 2:**
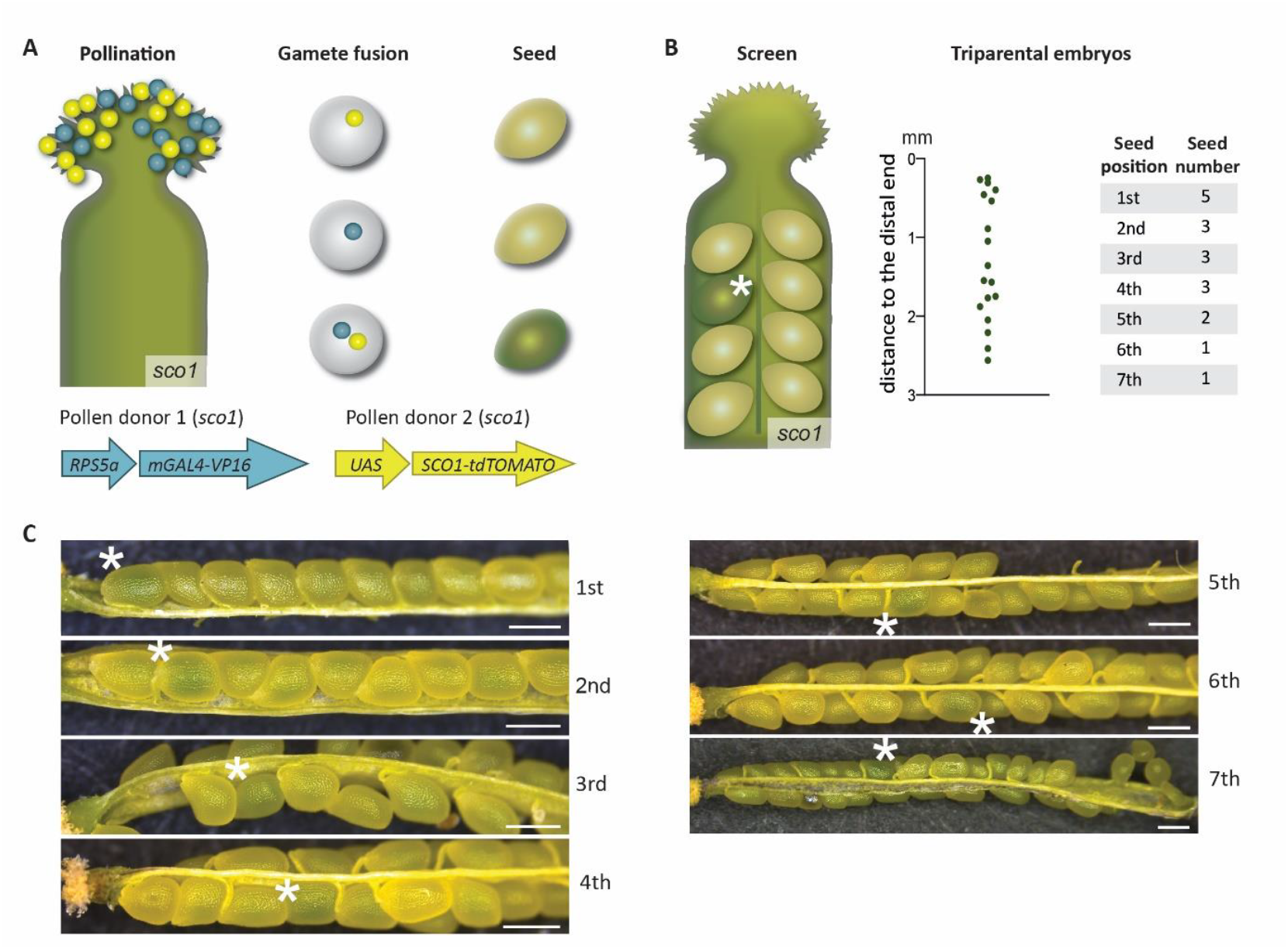
Proximity to the stigma positively affects the likeliness of polyspermy-induced triparental embryos. (A) In the HIPODS_*SCO1*_ *assay*, homozygous *sco1* mutants are pollinated with pollen from pollen donor 1 and pollen donor 2. Regular monospermy results in pale green seeds, while polyspermy involving both fathers activates the functional *SCO1* copy thereby restoring the darker green seed color. (B) Illustration showing an opened silique subjected to HIPOD_*SCO1*_ segregating a dark green seed indicative of a polyspermy-derived triparental embryo (asterisk). The distance of the rescued seeds and the seed position within the locule, both relative to the distal end of the style and measured from the funiculus are given. (C) Triparental embryo-segregating siliques 7 days after pollination (DAP). Seeds containing triparental embryos are marked by an asterisk and the position within the locule is indicated to the right. Scale bar: 0.5 mm. Figure adapted from Mao *et al*., 2020.

We dissected siliques 7 days after pollination (DAP) and determined the position of darker green seeds within the siliques from more than 100,000 analyzed seeds (Figure 2B). On average, there were approximately 34 seeds per *sco1* silique 7 DAP (34.36±8.77, n=36). In total, 30 polyspermy-recovered green seeds were identified. Of the seeds with clearly assigned positions, 18 were located within the first 7 seeds when counted from the distal end of the silique. The distance of the 18 seeds to the distal end of the style was determined and shown to range from 0.25 to 2.56 mm (Figure 2B, C). The average 7 DAP silique length of the *sco1* mutant varied from 7 to 8 mm. From this, we conclude that polyspermy events often occur at the distal end of the carpel where the ovules are more likely to receive more pollen tubes.

### Genetic component: Polyspermy is accession dependent

Given that polyspermy is a potent route towards polyploid plants, we asked whether polyspermy frequencies differ in plants from different accessions. For this, we introduced the HIPOD system into Arabidopsis plants of the Columbia (Col-0) accession and established homozygous lines for *pRPS5a:mGAL4-VP16* and *pUAS::BAR-YFP*, i.e. pollen donor 1 and pollen donor 2, respectively. We subjected Col-0 and L*er* plants to the HIPOD assay and observed a more than three-fold increase in polyspermy frequencies in the Col-0 compared to the L*er* accession (Table 1). The increased adaptive potential associated with whole genome duplications under stressful conditions^22-24^, along with our findings that the propensity to polyspermy varies among different plant accessions, indicates that susceptibility to polyspermy is an adaptive trait.

**Table 1:** Polyspermy frequencies are accession dependent. Table showing polyspermy frequencies obtained on intra- and inter-accession crosses using L*er* and Col-0.

| Female | Pollen donors | TP | Seed number | Psf* (%) |
| --- | --- | --- | --- | --- |
| Ler | Ler | 9 | 107376 | 0.020 |
| Col-0 |  | 19 | 92680 | 0.047 |
| Ler | Col-0 | 7 | 115096 | 0.013 |
| Col-0 |  | 32 | 95051 | 0.068 |

\*Polyspermy frequency (Psf) = # Triparental plants (TP) x 2 / total seed number x 100

Next, we aimed to determine whether the source of the different polyspermy frequencies between L*er* and Col-0 can be traced back to the male or female interaction partner. Reciprocal crossings between pollen donors and pollen acceptors from these accessions were performed. In addition, emasculated flowers from wild-type L*er* and Col-0 were crossed independently to pollen donors from both accessions. Interestingly, when pollinating the emasculated flowers from both accessions using L*er* pollen donors, Col-0 plants displayed 2.35 times higher polyspermy frequency when compared to L*er* plants (0.047 vs. 0.020; Table 1). When Col-0 pollen donors were used, Col-0 plants showed 5.23 times higher polyspermy frequency compared to L*er* plants (0.068 vs. 0.013; Table 1).

These findings indicate that both partners affect polyspermy ratios, however that the major differences between L*er* and Col-0 resides in the female. The causes are likely diverse, as the interaction between male and female components is complex and occurs across different spatial stages. This process starts with the interaction between pollen and the stigmatic surface, followed by regulatory mechanisms that govern pollen tube growth, sperm delivery, and gamete fusion, and continues through to variations in seed development and germination^25^. Against this background and, the low frequency of polyspermy, and the many potential polyspermy-influencing genetic differences between the two accessions, we estimate that screening several million plants would be required to identify the specific locus. Therefore, we decided not to pursue this approach.

### Three-parent plant breeding is applicable in crop plants

Given the tremendous potential of polyspermy-induced triparentage for speeding up breeding processes and bypassing hybridization barriers, we aimed to transfer the three-parent breeding technology from a laboratory setting to in-field crop plants. For this, we developed a new screening method for the sugar beet crop. This method involves a pollen donor (father 1) carrying a gene conferring sulfonylurea (SU) tolerance^26^, and a dominant red hypocotyl marker provided by the second pollen donor (father 2)^27,28^. Triparental seedlings derived from polyspermy can hence be discriminated against biparental ones via the combined occurrence of herbicide resistance and a red hypocotyl (Fig. 3A). We conducted a field trial in Italy during the 2020 growing season. To maximize stigmatic pollen density through wind pollination by the fathers and to provide balanced distributions of father ratios irrespective of wind direction, plants were sown in rows with alternating genotypes (Fig. 3B). Male-sterile, SU-sensitive mother plants with a green hypocotyl were used to collect seed derived the pollination experiment (Fig. 3A). After the growing season, seeds were harvested, dried, pre-processed, and sown in a greenhouse (Fig. 3C). The resulting offspring was treated with SU herbicide and herbicide-resistant plants were screened for red hypocotyl pigmentation. Herbicide-sensitive plants with a red hypocotyl were still able to germinate but either arrested in the two-leaf seedling stage or exhibited stunted and delayed growth. SU-tolerant plants with a green hypocotyl, indicative of biparentage were removed manually (Fig. 3C).

**Figure 3:**
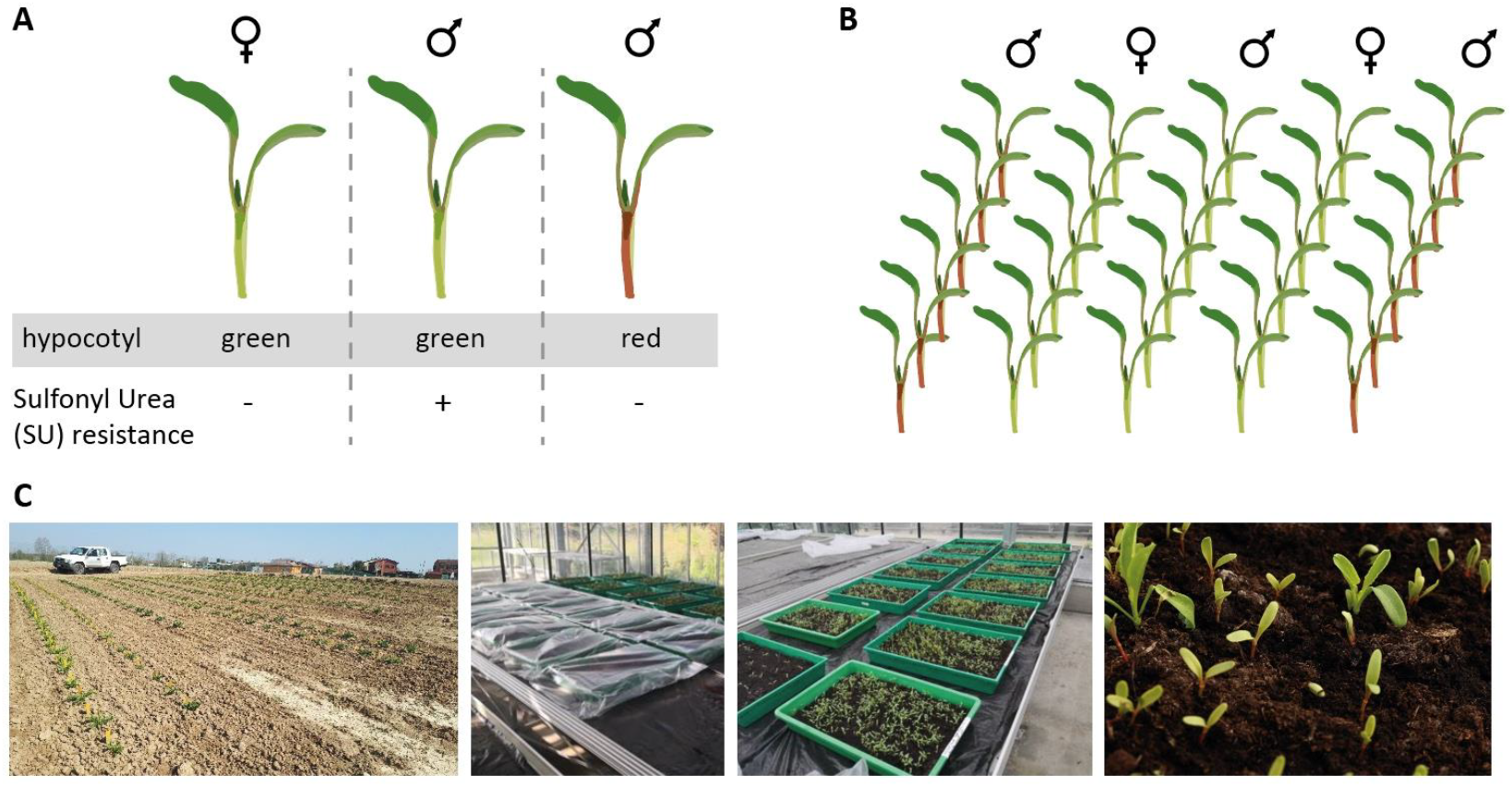
Experimental set-up for triparental sugar beet identification and field trial evaluation. (A) Illustration of the crosses involving two genetically distinct fathers. (B) Planting strategy: The mother plant and the two distinct fathers were arranged in adjacent rows to optimize stigmatic pollen density and balanced distribution of paternal pollen through wind pollination. (C)Field trial setup (left), seed sowing, and greenhouse cultivation (middle). After treatment with SU herbicide, most red hypocotyl plants remained at the cotyledon stage, while green hypocotyl plants grew normally (right).

Plants that exhibited a red hypocotyl and developed true leaves were subjected to ploidy analysis. While most tested plants exhibited a diploid profile, we identified 5 plants that showed a shift in the ploidy profile indicative of a triploid sugar beet (Fig 4A). These plants were repotted and sampled for DNA isolation (Fig. 4B). Subsequent PCR analysis and Sanger sequencing revealed a single-nucleotide polymorphism (SNP) conferring SU tolerance inherited from father 1 (Fig. 4C). Additionally, a 5 bp deletion conferring dominant red hypocotyl pigmentation from father 2 was confirmed (Fig. 4D). These results show that all plants segregated both selection markers, demonstrating that genetic material from two fathers was transmitted to a single egg cell.

**Figure 4:**
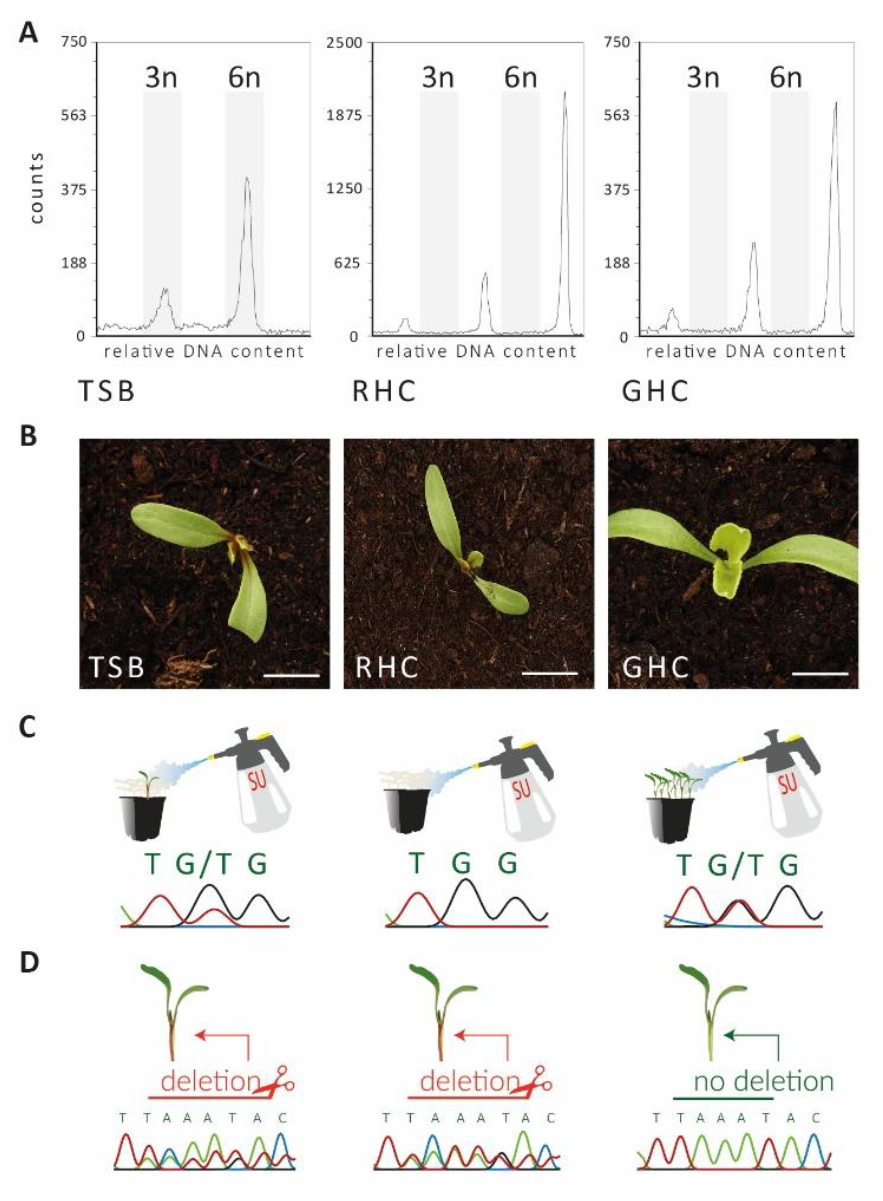
Triparental sugar beet selection. (A) Flow cytometric analysis of an herbicide-resistant plant with a red hypocotyl (left) reveals a distinct triploid profile compared to diploid wild-type controls with red (middle) and green hypocotyls (right). (B) Triparental candidate recovered from herbicide treatment (left), diploid wild-type offspring displaying red (middle) and green hypocotyls (right). (C) Sanger sequencing of the sulfonyl-resistance locus in the triparental candidate shows the presence of a G-to-T substitution at a 1:2 ratio, indicating one genomic copy from the herbicide-resistant father (left). The untreated red hypocotyl control shows no substitution (middle), while the green hypocotyl offspring has a 1:1 ratio (right). (D) Sequencing of the triparental candidate identifies a deletion that dominantly confers red hypocotyl color at a 1:2 ratio, suggesting one genomic copy from the red hypocotyl father (left). The red hypocotyl diploid offspring exhibits a 1:1 ratio (middle), and the green hypocotyl control lacks the deletion (right). TSB - Triparental Sugar Beet, RHC - Red Hypocotyl, GHC - Green Hypocotyl.

This study demonstrates that the generation of triparental plants depends on ecotype and stigmatic pollen density. By designing a wind pollination strategy that optimize equal paternal pollen density in sugar beet, we have transitioned three-parent crosses from a laboratory setting to field production and generated a triparental crop plant by instantly combining the genomes of one mother and two fathers. Our findings not only highlight the crucial role of polyspermy as a viable pathway for producing polyploid plants, but also demonstrate the practical applicability of three-parent breeding in crops when carefully designed and executed. This introduces a promising new tool for plant breeding. This innovation is particularly timely, given that the current pace of crop improvement is insufficient to meet the growing demands of a rapidly expanding global population. Several key publications have underscored the urgent need for an agricultural revolution to overcome this challenge. Despite extensive breeding efforts, elite cultivars often lack the climate-resilient traits that have naturally evolved in many wild varieties in response to environmental stresses^29^. Thus, finding innovative ways to introduce novel genetic traits into modern cropping systems is imperative^30^. However, breeding of these improved varieties is hindered by hybridization barriers that prevent crossing with stress-tolerant or pest-resistant wild varieties^29,31,32^. A potential solution lies in three-parent breeding technology, which has the potential to bypass the post-fertilization barriers of the endosperm by enabling selective polyspermy in the egg cell. This facilitates the introgression of previously incompatible wild varieties, contributing to the development of climate-resilient cultivars. The implications of this methodology are far-reaching, offering a transformative approach to crop breeding and reshaping our understanding of evolutionary biology. Particularly because this breeding approach avoids genetic engineering and requires minimal technical expertise and infrastructure, it is highly accessible to a wide range of users, fostering inclusivity. Let’s join forces to enable the development of resilient and sustainable crops for tomorrow.

## Material and Methods

### Plant growth conditions

Plants were sown in rows with alternating genotypes for in-field wind pollination during the 2020 growing season. Following pollination, sugar beet seeds were harvested, dried and pre-processed before being sown on trays with a 50:50 mix of P and T soil (supplier) and 6 liters of sand per 50 liters of soil. Trays were filled with 8 cm of loose soil, seeds were densely sprinkled and covered with 1-2 cm of soil. Each tray received an initial irrigation of 1.4 liters. Herbicide MaisTer® power was mixed with water and applied according to the manufacturer’s instructions. Trays were covered with plastic bags until germination, then opened gradually over two days to acclimatize seedlings before removing the covers. Cultivation took place in a greenhouse with temperatures ranging from 15 to 46°C, without controlling humidity, CO2, or temperature. Plants were additional treated with the fungicide Ortiva® (670 µl per tray) 10 days after germination.

Arabidopsis seeds were sown on the same soil containing pots and were subjected to cold treatment (4 °C) for 3 days in a Thermo Scientific Forma cold chamber. Plants germinated in a Conviron MTPS growth chamber under long day conditions (16 h light/8 h dark) at 23 °C and stayed under these conditions until bolting. Afterwards they were transferred to 18 °C Conviron MTPS growth chamber under long day conditions. For HIPOD_SCO1_, plants were kept at 23 °C after bolting.

### Emasculation and pollination

Three closed mature buds from the primary branch were emasculated using forceps. After 3 days, the stigma of the flower were double-pollinated using paint brushes containing pollen grains harvested separately from plants harboring *pRPS5a:mGAL4-VP16/+* and *pUAS::BAR-YFP/+* as described in Nakel *et al*. 2017. Limited pollination was carried out in a similar way with the help of an eyelash depositing approximately 15 pollen from each father. The HIPOD_SCO1_ experiment was carried out following the previous publication of Mao *et al*. 2020. The *sco1/-* mutant was emasculated and pollinated by using pollen grains taken individually from plants of *sco1/-pRPS5a:mGAL4-VP16/+* and *sco1/-pUAS::SCO1_tdTOMATO/+*.

### HIPOD screening and polyspermy frequency estimation

HIPOD screening and polyspermy frequency estimation was performed according to Nakel *et al*., 2017 and Mao *et al*., 2020. In short, images of seeds from double pollination were captured on a white background using a Canon EOS 700D camera. Seed numbers were estimated using an image-based recognition software (Count_seeds.py). To identify herbicide-resistant plants, seedlings were sprayed with BAYER Herbicide BASTA solution (1:1000 dilution) 2-3 times during the course of 15 days in equal intervals. For YFP expression analysis, sepals from open flowers were placed on a glass slide containing 10% (v/v) glycerol and images were captured using a Leica DMI6000b epifluorescence inverted microscope, equipped with a YFP ET, k (material number 11504165) and a DAPI ET, k (material number 11504203) filter cube. In the HIPOD_SCO1_ experiment, seed color screening was performed on dissected 7 days after pollination (DAP) siliques using the Leica S6 E or S8 APO stereomicroscope (Leica, Germany). Seeds with a color change were analyzed for tdTOMATO fluorescence and embryo and endosperm chromosome number.

For ploidy analysis, 0.5 cm^2^ leaf material (Sugar beet or Arabidopsis) was chopped with a razor blade in CyStain^®^ UV Precise P Nuclei Extraction Buffer (Sysmex Europe GmbH, Germany). The homogenate was then incubated with CyStain® UV Precise P Staining Buffer for 1 minute and filtrated through a 30 µM pore size CellTrics^TM^ nylon filter (Sysmex Europe GmbH). The flow through was analyzed using the CyFlow Ploidy Analyzer (Sysmex Europe GmbH).

Polyspermy frequency was estimated based on triparental plant formation. Only plants exhibiting herbicide resistance, a triploid genome, YFP expression and both the paternal constructs were considered triparental. Since HIPOD at its best detects only 50% of all polyspermy events, the polyspermy frequency was calculated using the formula below:

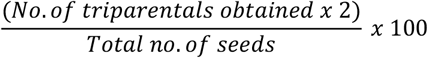

### DNA isolation, PCR and Sanger sequencing-based genotyping

Genomic DNA was isolated either using the MagMax™ Plant DNA Isolation Kit (Applied Biosystems) according to manufacturer’s instructions or using the Edwards method^33^. Plants were genotyped by PCR using primers 5′-TATGGGGTTTGGGCTACCAG-3′ and 5′-TCAATAAGGTTCTTCCATCA-3′ for amplification of the SU tolerance locus^26^, and 5′-GATGCTGGAACAGACACTAC-3′ and 5′-CTGTCACTTCATTGTCG-3′ for amplification of the red hypocotyl locus^27,28^. The resulting fragments were purified from a 1% Agarose Gel and extracted using NucleoSpin Gel and PCR Clean-up kit (Macherey Nagel) and analyzed via Sanger sequencing by LGC Genomics using the sequencing primers 5′-AGTTGCTCGACCAGATGCAGTG -3′ for SU tolerance and 5′-GGGTCATGGCAGAGTTAATTAGG-3′ for red hypocotyl locus.

## Acknowledgements

We are grateful to our team members for their discussions, helpful advice, and comments on the manuscript. We gratefully acknowledge the financial support from the European Research Council to R.G. (ERC Consolidator Grant “bi-BLOCK” ID. 646644, ERC Proof of Concept “TriVolve” ID. 957547).

## Author contributions

R.G. O.C., and T.N. conceived the study; S.J., Y.M., O.C., T.N. and R.G. designed research; S.J. performed the pollen density and plant accession studies; Y.M. performed the stigmatic proximity study; S.J., Y.M., T.N., D.T., T.B., and J.P. took part in HIPOD assays; O.C. performed the in-field wind pollination experiments of sugar beets; C.A.B. and T.N. performed the sugar beet seed counting, sowing, cultivation, triparental plant identification, and DNA fingerprinting with assistance from C.B.; R.G. wrote the paper with input from all other authors.

